# Embryonic origin of cancer in newborn twins

**DOI:** 10.64898/2026.05.10.722519

**Authors:** Barbara Walkowiak, Henry Lee-Six, Sadie Baskind, Manas Dave, Pooja Balasubramanian, Nathaniel Anderson, Jonathan Kennedy, Thomas R. W. Oliver, Rohan Verma, Sarita Depani, Simon Hannam, Tom Watson, Dyanne Rampling, Karin Straathof, Liina Palm, J. Ciaran Hutchinson, Olga Slater, Sam Behjati

**Author notes:** Equal contribution (Barbara Walkowiak, Henry Lee-Six, Sadie Baskind).

## Abstract

Studying monozygotic twins who present with identical cancers informs on the developmental origins of childhood tumours. Here, we performed whole genome sequencing on multiple tumour, normal, and placental samples to reconstruct the phylogeny of a soft tissue cancer that spread in utero between monozygotic twins. This generalisable and scalable approach allows us to dissect the earliest stages of twinning, revealing unexpected asymmetrical contributions of embryonic lineages to the placenta and each twin.

## MAIN

There are exceptional reports of identical twins (i.e. derived from the same zygote) who present with the same cancer^1,2^. Tumours in twins may arise independently, in particular in the context of a cancer-predisposing mutation in the germline^3^. Alternatively, they may result from *in utero* transfer of tumour cells from one twin to the other^4,5^. These scenarios may be distinguished by comparing the genetic changes in the two tumours: shared variants indicate a single origin followed by spread, while independent sets of mutations imply separate origins. Healthy cells also acquire somatic mutations over time, and so by looking at the pattern of sharing between the tumour and different normal samples, it is possible to reconstruct not only the life history of the tumour but also some of the earliest steps in development^6,7,8,9^.

Using this approach, we investigated the origin of lethal cancers in female monozygotic twins. A facial mass had been found in one girl (hereafter referred to as twin A) during fetal imaging, and at birth, tumour lesions were found to be disseminated both in her and her sister (twin B) (**Figure 1A-E, Supplementary Note 1**). While twin A had tumour deposits in multiple organs, twin B’s lesions were found in the brain and the skin. Biopsies of their tumours were reported as *MNF1*::*ZNF341* rearranged undifferentiated sarcomas (**Figure 1F, Supplementary Note 1**). Both twins, sadly, died soon after birth.

**Figure 1.**
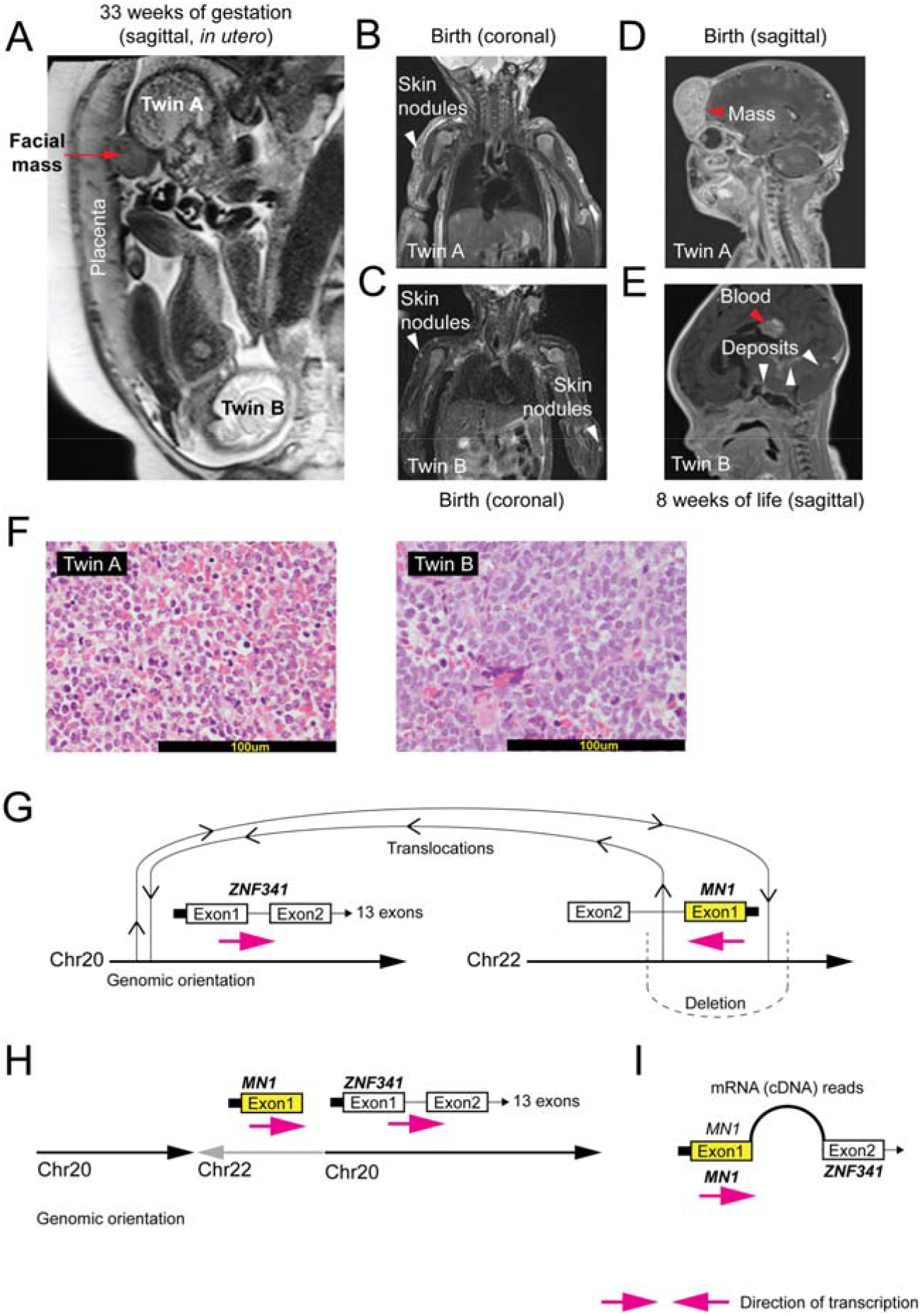
Presentation of twins with a sarcoma. **Panel A** shows a fetal MRI at 33 weeks gestation, one week before the twins were delivered. A facial mass was observed in twin A. **Panels B** and **C** show coronal MRI images of twins A and B respectively at birth. Both had skin nodules. **Panel D** shows the facial mass of twin A at birth. **Panel E** shows the intracranial haemorrhage (red arrow) noted in Twin B at 8 weeks of age, along with leptomeningeal tumour deposits (white arrows). **Panel F** represents haematoxylin and eosin stained sections of skin deposits in twin A and B, respectively, with identical small round blue cell morphology noted. **Panel G** presents the translocation that led to the formation of MN1::ZNF341 fusion. **Panel H** depicts the gene fusion that was found to be driving the tumours. **Panel I** shows the resulting cDNA derived from RNA sequencing.

We reconstructed the phylogeny of normal twinning, cancer formation and spread from normal and tumour tissues of both twins obtained at post mortem. We performed whole genome sequencing on 23 normal samples (six normal samples from each twin,and 11 placental samples) and 10 tumour samples (eight from twin A, two from twin B) (**Supplementary Table 1**). In addition, we sequenced 12 placenta samples enriched for trophoblast by laser capture microdissection (LCM). We called all classes of mutation and identified developmental somatic substitutions, i.e. mosaic variants that are shared between some cells across both twins and placenta. We additionally performed RNA sequencing to resolve and validate the *MN1::ZNF341* fusion (**Fig 1G-I, Extended Data Figure 1**).

We first examined the phylogeny of monozygotic twinning, which, in principle, can assume one of three configurations (**Fig 2A,B**). In our data, we identified three groups of mutations (**Fig 2C**). The first (**Fig 2C**, mutation A), was present in both twins. The second (**Fig 2C**, mutations B-H), was composed of mutations that made a major contribution to twin A. Some of these mutations approached a variant allele fraction (VAF) of 0.5 in twin A, indicating that the cell in which they arose gave rise to nigh all the cells of this twin. The third (**Fig 2C**, mutations I-T) group consisted of variants which were restricted to the tissues of twin B. Next, we assessed the contribution of twin lineages to the placenta, sampled systematically across six different territories (n = 18 samples; **Fig 2D-E, Extended Data Figure 2**). Unexpectedly, we found a dominant contribution of twin B’s lineages to placental samples.

**Figure 2.**
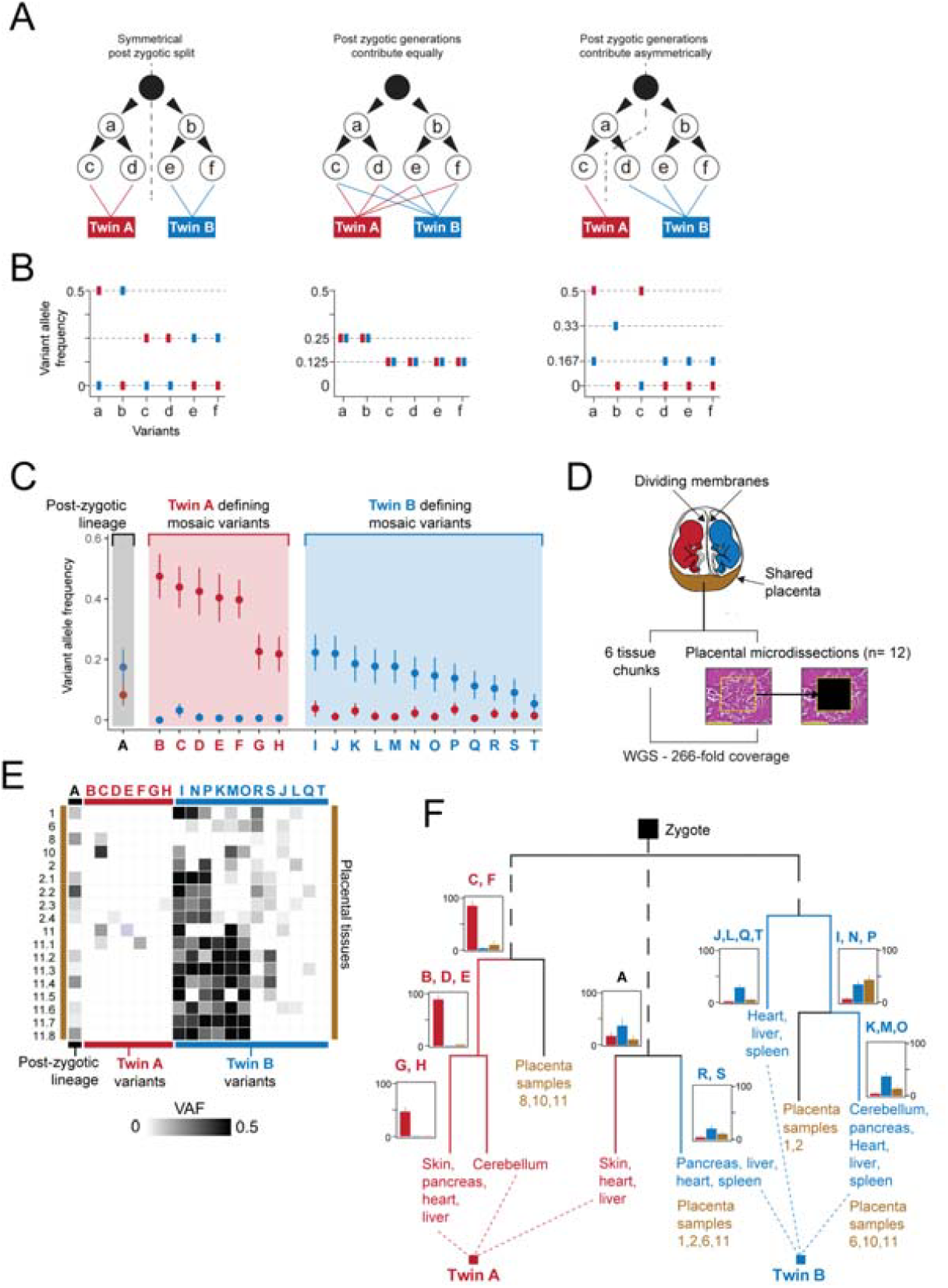
Reconstruction of early development. **Panel A** presents three different possible scenarios of the early events associated with twinning. **Panel B** presents the expected variant allele frequencies (VAFs) of early embryonic mutations under each scenario. **Panel C** presents VAFs of early embryonic mutations in normal samples of twin A and twin B. Error bars show 95% confidence intervals. VAF was calculated by aggregating reads from all normal samples, excluding the spleen sample from twin A and skin sample from twin B due to cross-contamination of those samples with material from the other twin. **Panel D** presents the study design to sample the placenta. **Panel E** compares the VAFs of early embryonic mutations in each of the bulk and LCM placental samples. **Panel F** presents the reconstructed phylogeny which shows a possible and congruent record of the early events associated with twinning.

From the distribution of developmental mutations across tissues we derived a phylogeny of twinning (**Fig 2F**) which illustrated the asymmetric fate of the earliest lineages: while one lineage (marked by mutations C and F in Fig 2F) contributed almost exclusively to twin A, another (marked by mutation A in Fig 2F) contributed to both twins and the placenta, and the third (marked by mutations J,L,Q,T,I,N,P in Fig 2F) contributed to twin B and the placenta (**Extended Data Figure 2, 3; Supplementary Note 2**). The sum of VAFs across the three parallel lineages was 0.5 in each twin, indicating that we have captured all early lineages that contributed to sampled tissues in both twins (**Extended Data Figure 4**).

We next reconstructed the origin of the sarcoma, distinguishing between two scenarios (**Fig 3A**): two independent tumours, or the origin of a tumour in one twin followed by transfer to the other. We exclusively found the genotype of twin A in tumour tissues, irrespective of whether they were sampled in twin A or in twin B, which establishes that the cancer formed in twin A and then transferred to twin B (**Fig 3B, Extended Data Figure 5**).

**Figure 3.**
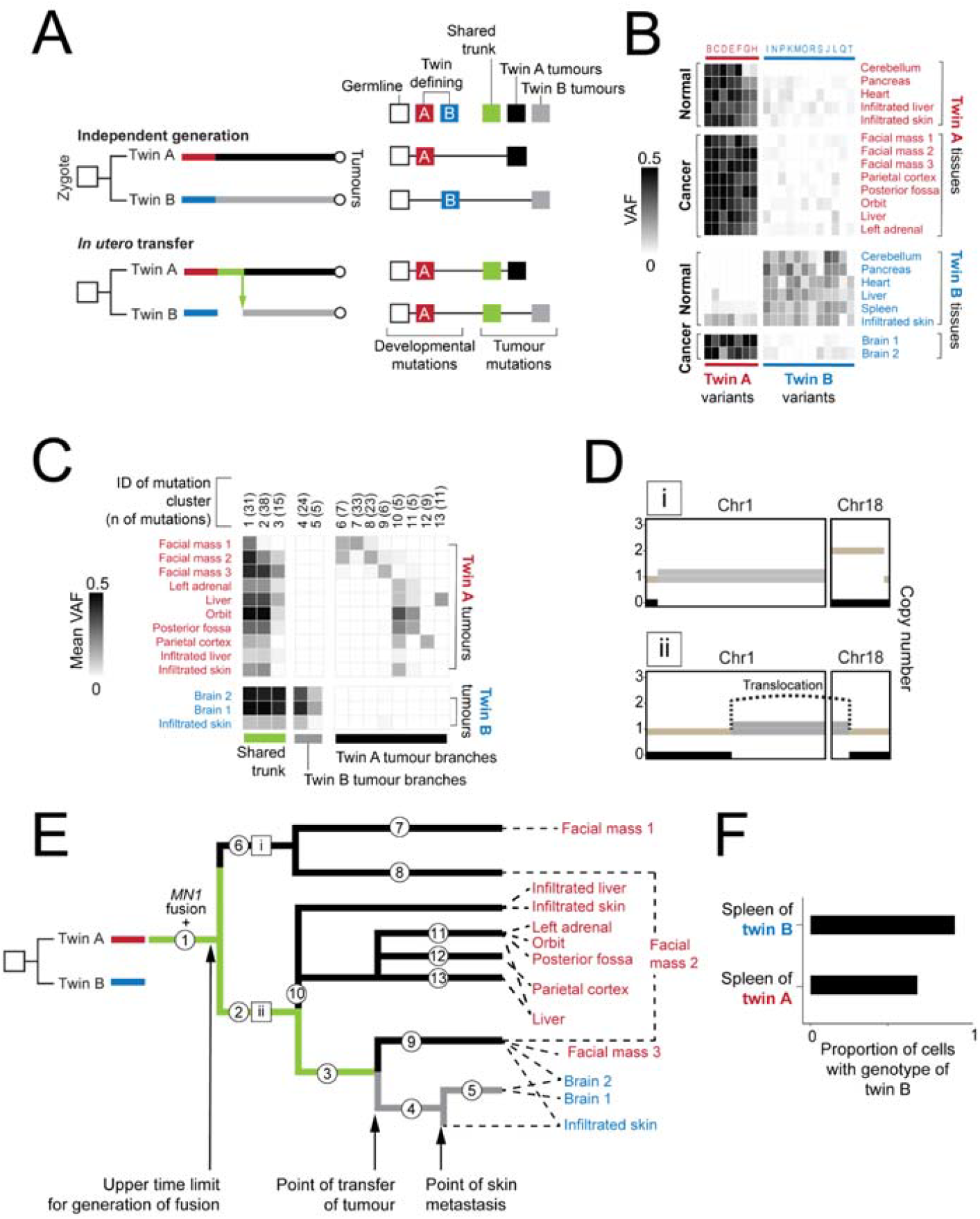
Evidence for tumour transmission and transfusion between twins. **Panel A** represents possible explanations for the presence of the tumour in both twins and the expected pattern of mutation sharing in the tumour in each case. **Panel B** shows VAFs of each early embryonic mutation (A-T) genotyped in normal samples of twins and tumour samples from twin A and twin B. **Panel C** shows the VAFs of mutation clusters 1-13 in each tumour and normal sample (‘placenta’ here represents aggregated bulk placental samples). **Panel D** shows the configuration of chromosome 1 and 18 copy number changes in two parallel clones of the tumour. The rearrangement between chromosomes 1 and 18 is indicated with a curved line. Phased chromosomes are indicated in beige and black. Segments of the chromosome that could not be phased are indicated in grey. **Panel E presents** the reconstructed phylogeny of the tumour, with the shared trunk highlighted in green. **Panel F** presents the estimates of the fraction of cells from twin B in the spleen of each twin.

We reconstructed the subclonal composition of the tumour using patterns of shared mutations and copy number alterations (**Figure 3C-E**). We find that tumour dissemination in twin A likely represented the spread of multiple clones, whereas the tumours of twin B likely spread only once and was a relatively late event (**Fig. 3E, Supplementary Note 3**). Assuming a linear mutation rate, we can time the origin of the tumour as occurring at the end of the first trimester, and the spread between twins as occurring late in the second trimester (**Supplementary Note 4**).

In assessing all tissues for twin-defining mutations, we found mutations of twin B, the tumour recipient twin, in the spleen of twin A (tumour donor twin) (**Figure 3F, Extended Data Figure 6**). These mutations were at remarkably high VAFs, consistent with 75% of the cells in the spleen of twin A actually deriving from twin B. These observations are consistent with a mixing of blood cells between the twins, with flow from twin B to twin A.

Our phylogenetic reconstruction provides a detailed account of twinning and establishes the key events associated with cancer formation and spread between twins. We can unambiguously state that the cancer arose from cells of twin A and transferred to twin B. Based on both clinical (**Supplementary Note 1**) and genetic observations, the most likely site of tumour origin was the facial mass in twin A. After early diversification of the tumour and multiple metastases within twin A, the tumour’s spread to twin B occurred late, and, most likely, only once. Taken together, these observations suggest that the spread of the tumour was a rare event and therefore not a foregone conclusion.

Our analysis resolves the clonal architecture of twinning and the contribution of the early cell lineages to both twins and the placenta in this case. Whilst there have been previous efforts to study twinning from post-zygotic mutations^12^, the clarity afforded by deep whole genome sequencing of multiple somatic and placental tissues in our investigation revealed marked lineage asymmetries in early embryogenesis. Both twins were mainly formed from separate early lineages, with only one mutation found in both twins. Remarkably, with the caveat that we could not sample the entire organ, the placenta was mostly derived from cells that contributed to twin B. Overall, as well as mapping tumour spread *in utero*, our work delineates a powerful approach to reconstructing the earliest steps that lead from the zygote to the genetic segregation of twins.

## ACKNOWLEDGEMENTS

This research was funded by the Wellcome Trust (institutional grants [206194, 220540/Z/20/A] and personal fellowship to S.B. [223135/Z/21/Z]). H.L.S is a recipient of a research fellowship from Trinity College, Cambridge. This research was supported by the NIHR GOSH Biomedical Research Centre and NIHR Cambridge Biomedical Research Centre (NIHR203312). The views expressed are those of the authors and not necessarily those of the NHS, NIHR or Department of Health. We are indebted to this family for participating in our research.

## METHODS

### Ethics statement

The children’s parent provided written informed consent for participation in our research study which has been approved by a National Health Service Research Ethics Committee (reference 16/EE/0394).

### Data availability

All sequencing data have been deposited in the European Genome-phenome Archive (accession number in progress).

### DNA extraction and sequencing

Bulk DNA was extracted from fresh frozen tissues using AllPrep DNA/RNA minikit (Qiagen 80204). Placental samples were fixed in formalin and embedded in paraffin. They were extracted using the Qiagen QIAamp DNA FFPE Tissue Kit according to the manufacturer’s instructions. Short-insert genomic libraries were constructed and 150 base pair paired-end sequencing reads were generated on the Illumina HiSeq X or Novaseq platforms according to Illumina no-PCR library protocols. Sections of the placental tissue were prepared for LCM according to a previously described protocol^1^. Sample metrics are shown in **Table S1**.

### RNA extraction and sequencing

RNA extraction and sequencing was performed as previously described^2^.

### DNA sequencing alignment

All DNA sequences were aligned to the GRCh38 reference genome using the Burrows-Wheeler (BWA-MEM) algorithm^3^.

### RNA sequencing alignment

RNA sequences were aligned to the GRCh37 reference genome and aligned using STAR^4^. Reads that identify the MN1-ZNF341 fusion are shown in **Table S2**.

### Somatic mutation calling

We called mutations from different classes using the pipeline established at the Wellcome Sanger Institute. Substitutions were called with the CaVEMan algorithm^5^ against an *in silico* generated normal GRCh38 reference genome. We removed variants with reads with median mapping quality below 140 (ASMD>=140) and required that more than half of the reads supporting variants were not clipped (CLPM=0). For all reads, we set a cut-off base quality (25) and mapping quality (30). We generated a pileup from all .bam files. Small insertions and deletions (indels) were identified using the Pindel algorithm^6^. Chromosomal rearrangements were identified with the BRASS algorithm (https://github.com/cancerit/BRASS)^7^ (**Table S3**). Copy number alterations were identified using ASCAT^8^ (**Table S4**).

### Somatic mutation filtering

We employed a series of filters to remove likely artefactual mutations, and required a mutation to fulfil the following criteria to call it somatic:

⍰ Position mapped to chromosomes 1-22 or chromosome X (not chromosome Y)
⍰ No indels present within 30 base pairs of the substitution
⍰ Minimum 300 reads spanning the position across normal samples from twin A and twin B
⍰ Maximum 600 reads spanning the position across normal samples from twin A and twin B
⍰ Presence of minimum 4 variant-reporting reads and VAF minimum 0.1 in at least one sample
⍰ No significant strand bias observed in reads from all samples from twin A and twin B (normal and tumour), tested with Fisher’s exact test (alpha 0.05, corrected for multiple testing using Bonferroni correction).
⍰ Ratio of the number of reads identified in high-quality (mapping quality 30) to low-quality (mapping quality 5) minimum 0.7 across all samples from twin A and twin B (normal and tumour). This filter removes variants on poorly mapped reads, which could represent artefacts and for which we may not be able to reliably estimate the VAF.

Mutations retained and excluded after each filter are shown on **Supplementary Figure 1**.

### Germline mutation filtering

To remove germline variants, we perform a one-sided exact binomial test to identify variants which are heterozygous in all cells (VAF > 0.5) as described in Lawson et al.^9^. Briefly, for each twin, we aggregated reads across all normal samples and obtained the VAF of each mutation by dividing the number of variant-reporting reads by the total number of reads and tested if this mutation is likely to be germline (VAF > 0.5) in this twin (at alpha 0.01). To be excluded, the mutation has to be classified as germline in both twins. P-values were corrected for multiple testing with Bonferroni correction. 1,069 mutations were retained after excluding artefacts and likely germline mutations identified in this way. In this set of mutations, we identified 617 mutations present in all normal samples and, based on their VAF ∼0.3 and elevated coverage, found that those are more likely variants associated with copy number-altered regions in the germline (**Supplementary Figure 2**). We excluded those variants by excluding mutations with coverage (aggregated over all normal samples) above 1.25 or below 0.75 median of the depth of all mutations present on the chromosome, and mutations which were present in clustered (defined as 10 mutations of the 1,069 being present within a region of 50 kb). After excluding sequencing artefacts and likely germline mutations, we obtained a set of 542 mutations. We manually reviewed all of these mutations on Jbrowse^10^, checking for the mutation occurrence in both read orientations, at different read positions and presence on reads that with high sequencing and mapping quality and were not affected by the presence of a nearby insertion or deletion that could lead to mis-estimation of the mutation VAF. In addition, we excluded variants which, following this filtering process, could not be excluded to be germline. Following manual quality control, we retained 254 mutations that were used to reconstruct the phylogeny. Mutations identified as germline were removed.

### Genotyping of placental samples

Substitutions in LCM placental samples were called with the CaVEMan algorithm^5^ against an in silico generated normal GRCh38 reference genome. We required cut-off base quality (30) and mapping quality (30). We generated a pileup from all .bam files using the cpgVAF function, searching for the 20 early embryonic mutations and 212 tumour mutations. We then calculated the VAF for each of the mutations classified as early embryonic in each of the trophoblast samples, as well as all bulk placental samples aggregated together. Early embryonic and tumour mutations were genotyped in bulk placental samples using bcftools mpileup^11^ with cut-off base quality (30) and mapping quality (30).

### Estimation of tumour cell fraction in normal samples

Sharing of somatic mutations between tumour and normal samples could result from their shared developmental origin or infiltration of normal samples with tumour cells. To identify tumour-infiltrated samples, we used two approaches based on the presence of tumour-specific mutations and evidence of copy number alterations in normal samples. Results from both approaches were congruent, with estimates from copy number alterations being more sensitive to low-level tumour infiltration.

First, purity can be estimated by taking advantage of the loss of chromosomes 1q and 18p in tumour samples. Under the assumption that those alterations are specific to tumour cells and no copy number changes are present in normal samples, one would expect the VAFs of heterozygous SNPs and germline mutations in those regions to be at 0.5, while in a tissue sample which is infiltrated by tumour cells, we expect a systematic deviation from 0.5. We phased heterozygous SNPs on chr1q and chr18p; all SNVs in the copy-number altered regions which were found at VAF > 0.75 in posterior fossa tumour sample (purest tumour sample available) were assigned to one chromosome, while all SNVs present at VAF < 0.25 were assigned to the other chromosome. In each normal sample, we calculated the median VAF of mutations found on either chromosome and derived the expected fraction of cells which lost the allele (i.e., tumour cells). Purity estimates based on mutations on chr1q and chr18p were consistent for each sample. To obtain the final estimate of purity, we averaged the estimate based on chr1q SNPs and chr18p SNPs. The deviation from the expected value of 0.5 was consistent across the entire lost segment in both chr1q and chr18p for each sample (**Supplementary Figure 3**), arguing against focal copy number alterations in normal samples that would have confounded our estimates. Estimates are shown in **Table S5**.

To validate our estimates of purity, we also estimated tumour infiltration in normal samples by determining the fraction of reads reporting clonal tumour mutations in normal samples (**Supplementary Figure 4**). First, we identified 168 mutations present in all tumour samples and with VAF (aggregated across all tumour samples) min 0.2, and plotted VAF in the tumour against VAF of these mutations in each normal sample. For samples with no tumour infiltration, we expect to identify a cluster of mutations at VAF > 0.2 in the tumour and VAF = 0 in the sample. If the normal sample is heavily tumour-infiltrated, we expect that no separate cluster of mutations at VAF = 0 in normal will be present. With this line of reasoning, we identified highly pure samples and selected mutations which were present at VAF > 0.2 in the tumour, absent from highly pure samples, and at VAF < 0.1 in aggregated twin A samples to exclude potential embryonic mutations.

The observed VAF of a clonal tumour mutation each sample is given as

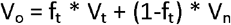

where:

⍰ V_o_: VAF observed in the sample
⍰ V_t_: VAF of the mutation in tumour cells
⍰ V_n_: VAF of the mutation in normal cells
⍰ f_t_: fraction of tumour cells
⍰ (1-f_t_): fraction of normal cells (1-f_t_)

As we searched for mutations that are clonal in the tumour and absent in normal samples, we assume V_t_ to be 0.5 and V_n_ to be 0. The tumour cell fraction is given as

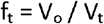

Estimates are shown in **Table S6**.

### Classification of somatic mutations and reconstruction of normal phylogeny

We categorized each of the mutations present in the set of 254 high-confidence mutations as belonging to one of the following categories:

⍰ present in normal samples from both twins
⍰ present in normal samples in twin A only
⍰ present in normal samples in twin B only
⍰ specific to the tumour (allowing for presence in tumour-infiltrated normal samples)
⍰ present in a single normal sample
⍰ present in a single tumour sample

A phylogeny was then constructed manually according to the pattern of mutation presence and VAF across samples. We also considered which trophoblast samples each normal mutation was confidently identified in. Assignment of mutations to the phylogeny is shown in **Table S7**. Coordinates of 20 early embryonic mutations used to reconstruct the early events associated with twinning (labelled A-T) are shown in **Table S8**. VAFs of the 20 early embryonic mutations used to reconstruct twinning in each sample are given in **Table S9**. VAFs of each of the tumour mutations are given in **Table S10**.

### Reconstruction of tumour phylogeny

We manually clustered 212 tumour-specific mutations into 13 clusters, each containing at least 5 mutations, based on the following criteria:

⌷ Cluster 1: mutations present in all tumour samples, VAF > 0.1 in the facial mass (sample 1)
⌷ Cluster 2: mutations present in min 7 tumour samples, absent from the facial mass (sample 1)
⌷ Cluster 3: mutations found in tumour samples from twin B, facial mass (sample 2 and 3), brain orbit, posterior fossa, left adrenal and liver tumour samples
⌷ Cluster 4: mutations identified only in tumour samples from twin B, VAF > 0.1 in both samples
⌷ Cluster 5: mutations identified only in tumour samples from twin B, VAF > 0.1 in one sample
⌷ Cluster 6: mutations found at VAF > 0.1 only in facial mass (sample 1 and 2)
⌷ Cluster 7: mutations found at VAF > 0.1 only in facial mass (sample 1) tumour
⌷ Cluster 8: mutations found at VAF > 0.1 only in facial mass (sample 2) tumour
⌷ Cluster 9: mutations found at VAF > 0.1 only in facial mass (sample 3) tumour
⌷ Cluster 10: mutations found at VAF > 0.1 in right parietal, brain orbit, posterior fossa, left adrenal and liver tumour samples
⌷ Cluster 11: mutations found at VAF > 0.1 in brain orbit, posterior fossa, left adrenal and liver tumour samples, but not in the right parietal tumour
⌷ Cluster 12: mutations found at VAF > 0.1 only in right parietal tumour sample
⌷ Cluster 13: mutations found at VAF > 0.1 only in liver tumour sample

We reconstructed the phylogeny of the tumour based on VAFs of mutations of each cluster in each sample, following the pigeonhole principle^7^. Assignment of each tumour mutation to a cluster is provided in **Table S11**. Mean VAFs of each cluster in each sample are given in **Table S12**.

### Estimation of cell transfer between twins

We estimate the levels of twin-to-twin transfusion using VAF values of twin A- and twin B-specific mutations. The observed VAF of a mutation in twin A spleen is given as

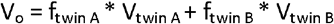

where:

⍰ V_o_: VAF observed in the spleen sample
⍰ V_twin A_: VAF of the mutation in twin A (non-spleen)
⍰ V_twin B_: VAF of the mutation in twin B (non-spleen)
⍰ f_twin A_: fraction of twin A cells in twin A spleen
⍰ f_twin B_: fraction of twin B cells in twin A spleen

For the 7 twin A-specific mutations, we estimate twin A VAF using non-spleen normal samples and assume VAF in twin B to be 0. We can calculate f_twin A_ as

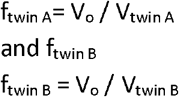

Estimates are provided in **Table S13**.

## Code availability

Custom code developed for the purpose of this analysis is available on GitHub under the link: https://github.com/BehjatiLab/tumour_transmission_mz_twins

## EXTENDED DATA FIGURES

**Extended Data Figure 1.**
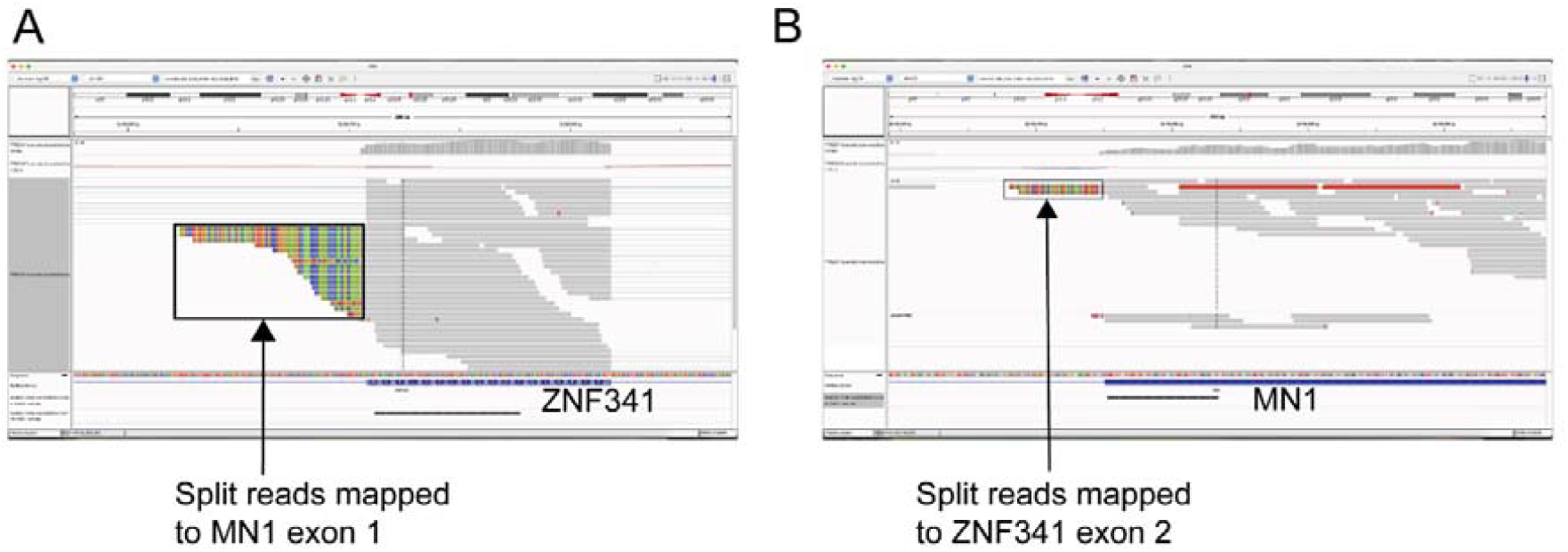
Reconstruction of the fusion from bulk RNA-seq reads from the twin A liver sample (contaminated with tumour). **Panel A** shows the view of RNA-seq reads of ZNF341, including split reads that map to MN1 exon 1. **Panel B** shows the view of RNA-seq reads of ZNF341, including split reads that map to ZNF341 exon 2.

**Extended Data Figure 2.**
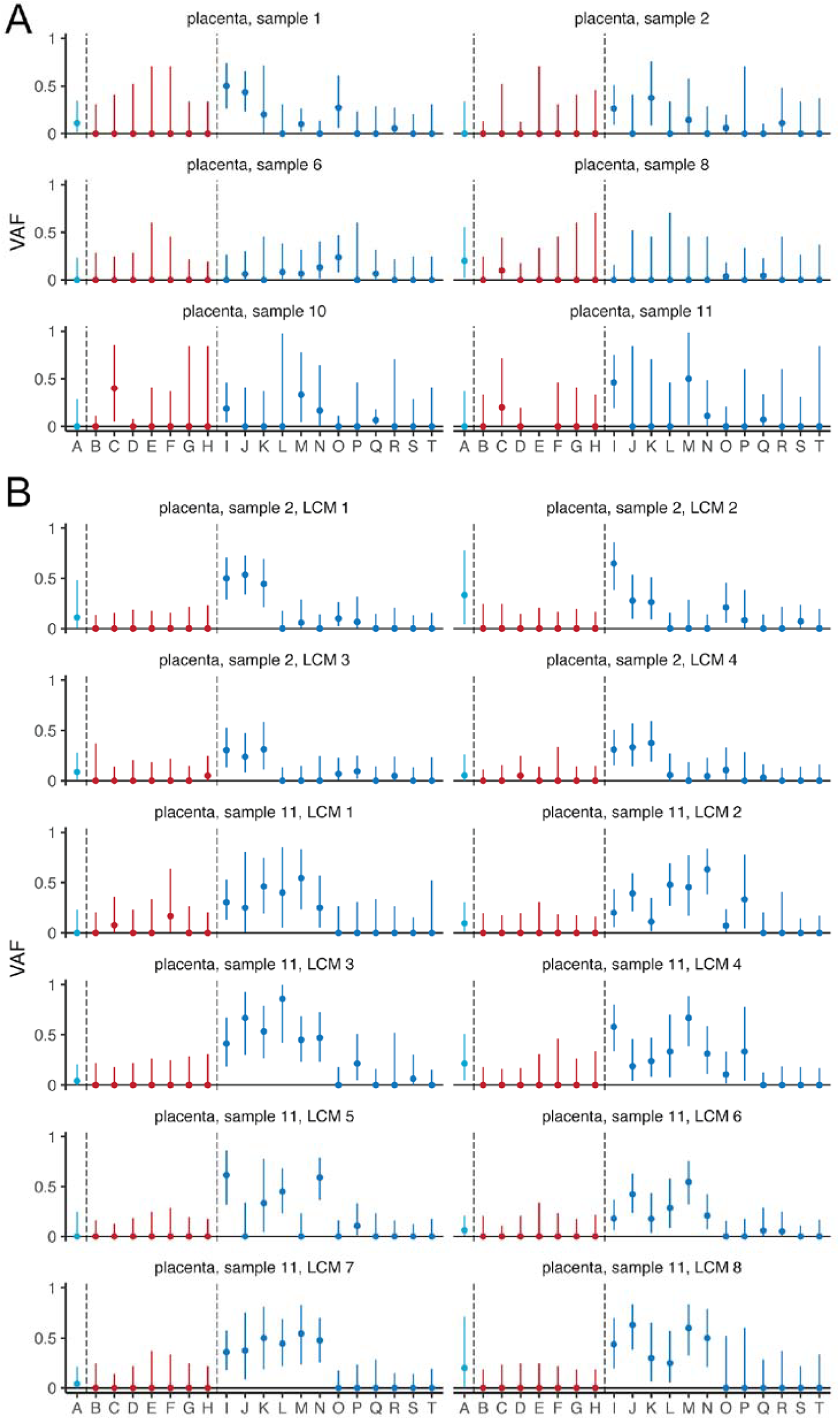
Estimates for variant allele frequencies (VAF) of the early embryonic mutations in each placental sample (A - bulk placental samples, B – LCM placental samples enriched for trophoblast). Error bars show 95% confidence intervals.

**Extended Data Figure 3.**
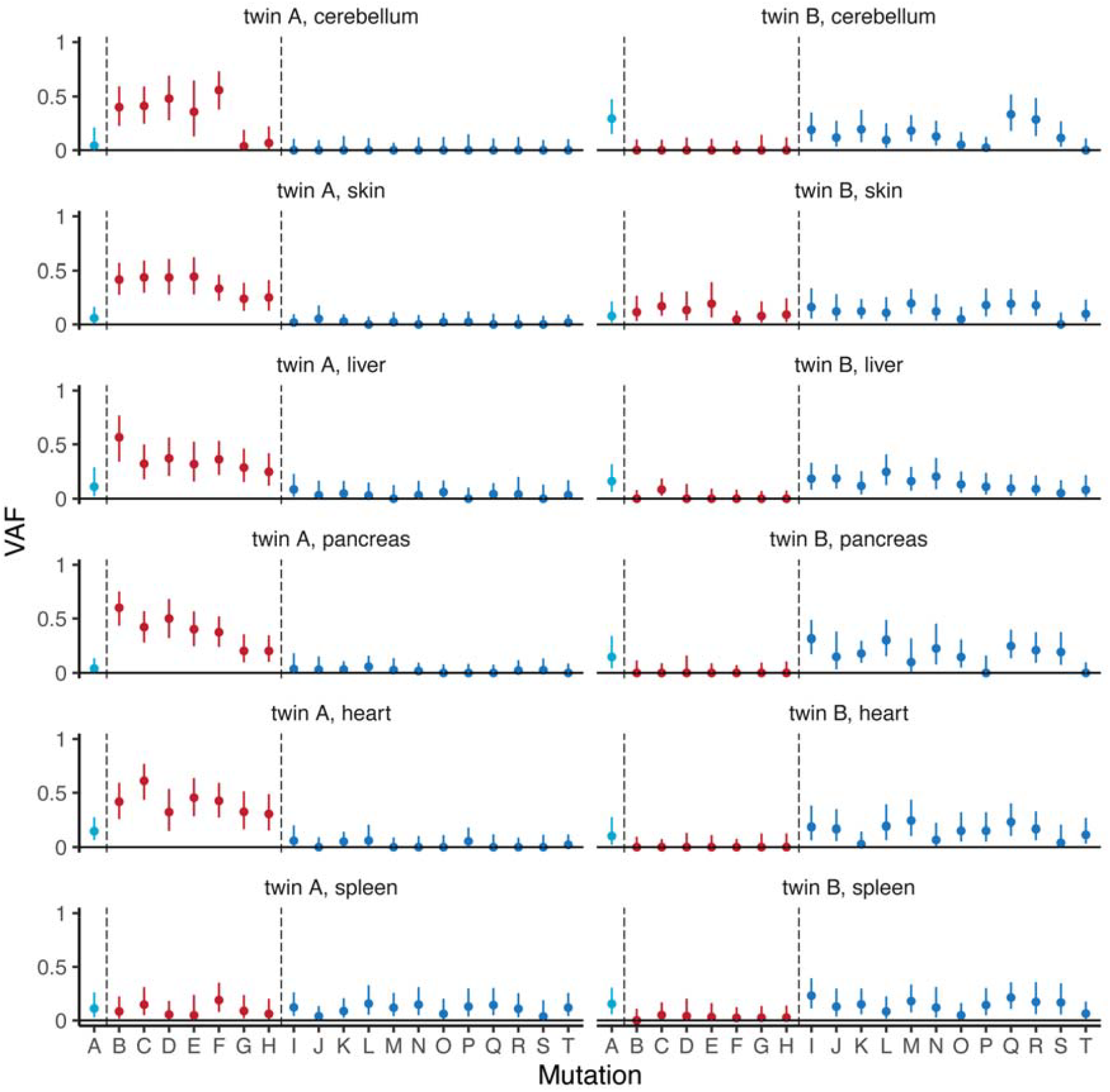
Estimates for variant allele frequencies (VAF) of the early embryonic mutations in each normal sample. Error bars show 95% confidence intervals.

**Extended Data Figure 4.**
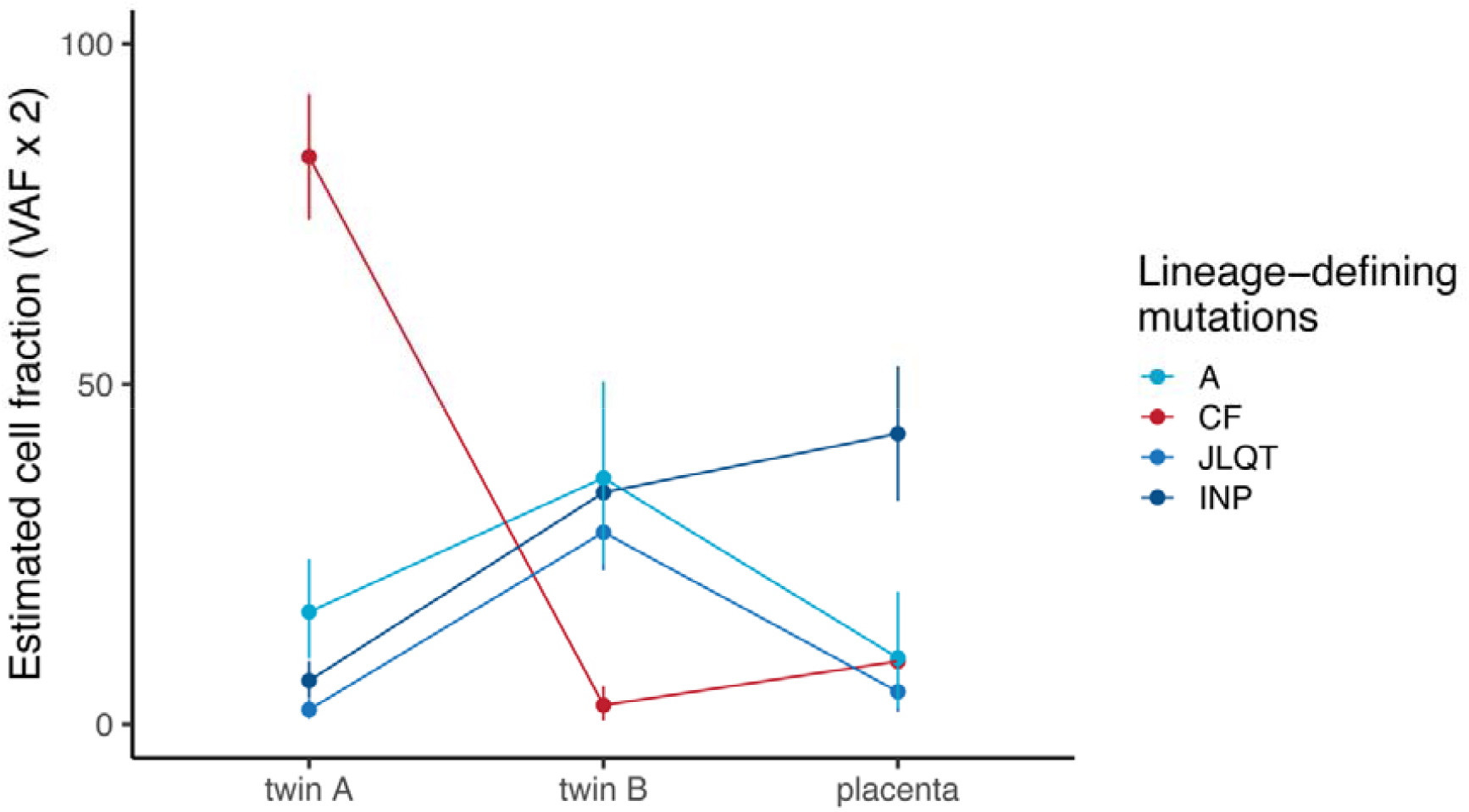
Estimates of the contribution of each major lineage marked by a group of mutations to each twin and the placenta. Error bars show 95% confidence intervals.

**Extended Data Figure 5.**
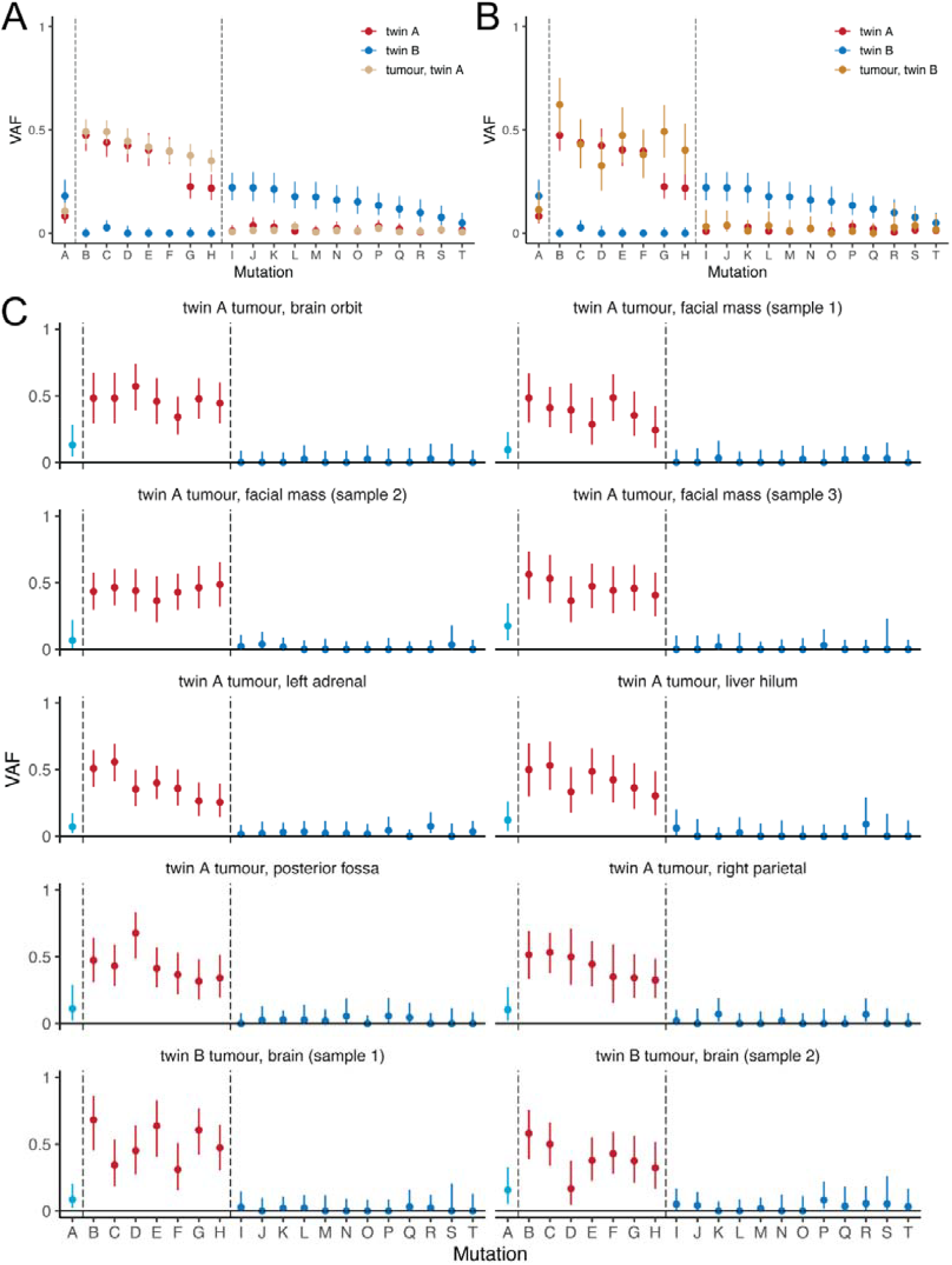
Estimates for variant allele frequencies (VAF) of the early embryonic mutations in the tumour. **Panels A**-**B** present VAFs of early embryonic mutations in twin (**A**) and twin B tumour (**B**) compared to variant allele frequencies across other tissues of each twin. Samples of spleen of twin A and skin of twin B are excluded from the aggregate values of normal samples due to contamination with mutations from the other twin. **Panel C** presents VAFs of early embryonic mutation in each tumour sample.

**Extended Data Figure 6.**
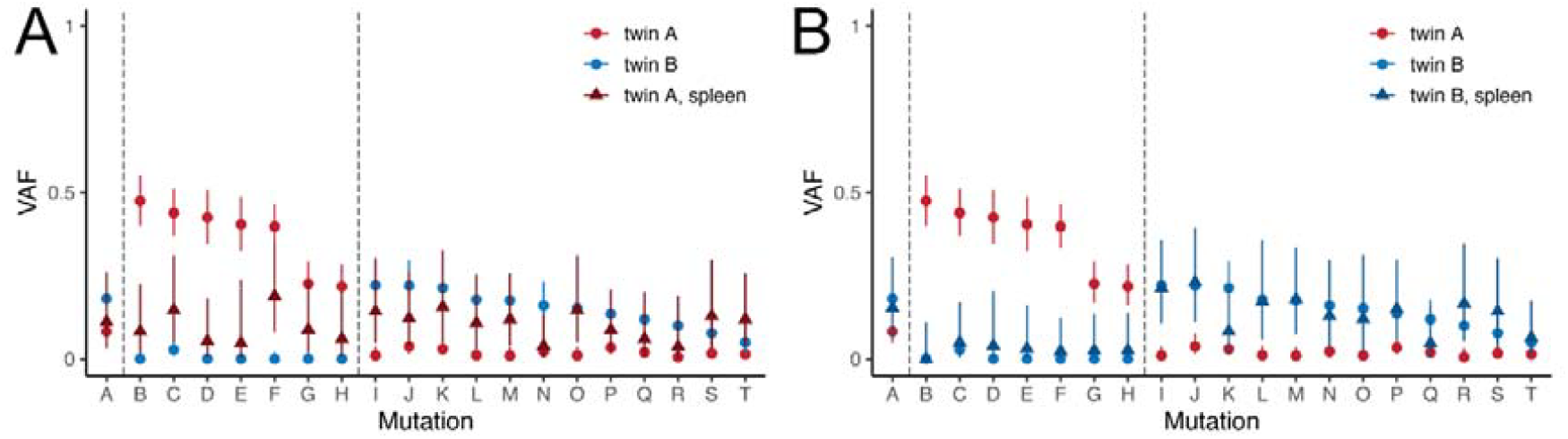
Estimation of twin-twin transfusion. **Panels A**-**B** show variant allele frequencies of early embryonic mutations in twin A spleen (**A**) and twin B spleen (**B**) compared to variant allele frequencies across other tissues of each twin.

