## Supplementary material for "Embryonic origin of cancer in newborn twins"

*Equal contribution

**Supplementary notes**

1. Histology of tumour samples
2. Cell transfer between twins
3. Clonal composition of the tumour in twin A and twin B
4. Timing of tumour origin in twin A and spread to twin B

**1 Histology of tumour samples**

Skin tumour deposits from both twins were reported as poorly circumscribed subcutaneous nodules composed of small round blue cells with brisk mitotic activity but not other specific morphological features. There was no neurofibrillary stroma and no cellular differentiation. Tumour samples from both twins were reviewed together and were noted to be morphologically identical. Immunostaining for Desmin, Myogenin, CD99, CD45, PHOX2B, and CD117 was negative. CD45, CD68 and CD168 were expressed in a few scattered intralesional cells, likely representing infiltrating T lymphocytes. INI-1 staining/ expression was retained in tumour cells. The neoplastic cells strongly and diffusely expressed CD56. Both tumours were later found to diffusely and strongly express SF1 in keeping with the characteristics published in the MN1 case series^12^.

Taken together, the features of the cancers in both children were of a small round cell tumour most consistent with an undifferentiated sarcoma. They were not consistent with neuroblastoma, leukaemia, mast cell disease, PNET, rhabdoid tumour, Wilms tumour or rhabdomyosarcoma.

**2 Cell transfer between twins**

We note that mutations which we term ‘twin-A specific’ were found at very low VAFs in the spleen and liver samples of twin B, and similarly, mutations called ‘twin B-specific’ were identified at very low VAFs in samples from twin A (**Extended Data Figure 2**). As monozygotic monochorionic twins share circulation *in utero*, those observations can be parsimoniously explained by blood transfer between twins.

Twin B specific mutations were observed at high levels in the spleen of twin A, and at low (VAF < 0.1) levels in the heart, pancreas, liver and skin. No twin B-specific mutations were found in the cerebellum. While we cannot formally exclude this being due to some low-level contribution of early cell lineages to twin A, we find blood transfer from twin B to be the more likely explanation for the following reasons:

1)	The elevation of those mutations in the spleen compared to other tissues argues for this being due to transfusion, especially in this clinical context where twin A presented with active extramedullary haematopoiesis.

2)	There is a moderate positive correlation between the VAF of twin B-specific mutations in twin B and spleen of twin A (Pearsons’ r 0.42), further arguing for this being due to blood transfer.

3)	Very low levels of twin B-specific mutations in other tissues of twin A are consistent with low-level tissue infiltration by blood cells (e.g., immune cells), expected in bulk samples

4)	The estimated contribution of lineages marked by mutations A, and mutations CF, to twin A is 0.999 (**Extended Data Figure 4**), indicating that we have likely captured all early cell lineages which formed twin A and are not missing other lineages forming this twin.

Twin A-specific mutations were observed in the spleen (6 / 7 twin A-specific mutations) and liver of twin B (1 / 7 mutations). The elevation of the VAF of those mutations specifically in the spleen is, again, congruent with this being a result of the sharing of the circulation between twins *in utero*. However, in this case, it is also possible that the sharing is due to the presence of the tumour, as microscopic tumour deposits have been identified in the spleen of twin B.

Given that our estimates suggest blood flow from twin A to twin B only occurred at low levels, it is not surprising that we do not see any evidence of twin A-specific mutations in other tissues.

**3 Clonal composition of the tumour in twin A and twin B**

The largest tumour mass in twin A was composed of multiple subclones (**Fig 3C-D, Supplementary Figure 5-6**), and metastases within twin A exhibited contributions from many of them, indicative of polyclonal dissemination. In contrast, the intracranial tumours and skin samples of twin B showed the presence of only two lineages, identified by mutations from clusters 4 and, at low levels, 9 (**Figure 3C, Supplementary Figure 6).** The similarity in clonal composition of these metastatic deposits in twin B suggests that the transfer of cells from twin A to twin B may have occurred only once. Mutations found in all samples of twin B’s tumour (cluster 4, **Figure 3C**) were clonal in her skin infiltrate but subclonal in her brain lesions (**Supplementary Figure 7**).

Tumour of twin B shows evidence of mutations which we have placed in clusters 1, 2, 3, 4, 5 and 9. Because clusters 4-5 and 9 are parallel and cannot be nested within each other, this indicates that the metastasis was not monoclonal (i.e., it was contributed by cells from at least two different lineages).

We considered whether there were one or more metastatic events from twin A to twin B. While we have not sampled the whole tumour of twin B, the data from samples available to us is consistent with a single metastatic event. In both the brain and the skin, we find evidence of the same lineages (marked by mutation clusters 1-5 and 9), with lower contributions of cluster 9 than 1-5 in both the brain and the skin. Achieving the same clonal composition in two independent metastases is unlikely, although it cannot be excluded.

Our data are consistent with tumour spread within twin B occurring from the brain to the skin. In both brain samples, the VAFs of mutations in cluster 4 (twin B-specific) are significantly lower than mutations in the cluster of truncal mutations (**Figure 3C**), while in the skin, the VAFs of mutations in those clusters do not significantly differ. If the metastasis occurred from the skin to the brain, the VAFs of those mutations would have been expected to be clonal also in the brain. However, it is important to note that we cannot exclude metastases from other regions of the skin, and only sampling the whole of the tumour would unambiguously resolve this question.

**4 Timing of tumour origin in twin A and spread to twin B**

We timed the transfer of the tumour from twin A to twin B by quantifying the mutations along the tumour phylogeny. The longest tumour branch was 120 substitutions, comprising 113 tumour variants and seven twin A-defining mutations (**Fig 3C-E, Supplementary table S11**). As 91 variants are shared by both twins’ tumours, the tumour spread at ~75% (91/120) of its mutational age (although we cannot exclude earlier spread due to undetectable subclonal mutations). To estimate when the cancer’s cell of origin ceased to be a normal cell, we timed the generation of the *MN1::ZNF341* driver fusion. As the fusion is clonal it must have occurred before the end of the tumour trunk, which is composed of 31 clonal substitutions and seven embryonic mutations. It occurred, therefore, during the first third of the tumour’s mutational lifespan ( (31+7)/120). Those estimates provide a relative timing of tumour origin and spread, although we note that we are unable to detect subclonal mutations present at a very low VAF, which may impact the accuracy of our estimates.

Mutational times may be converted into real times if we assume a linear mutation rate. This assumption is supported by two observations: first, the mutation burden of the tumour is low compared to other childhood cancers (**Supplementary Figure 8**); and second, the trinucleotide context of these substitutions is consistent with endogenous processes that are thought to act as cellular clocks (signatures 1 and 5)^11^ (**Supplementary Figure 9**). Accordingly, the gene fusion arose in the first trimester (38/120 variants × 40 weeks (the age of twin 2 at the time of sampling) = ~13 weeks) whilst tumour spread occurred during the second (91/120 variants × 40 weeks = ~26 weeks).

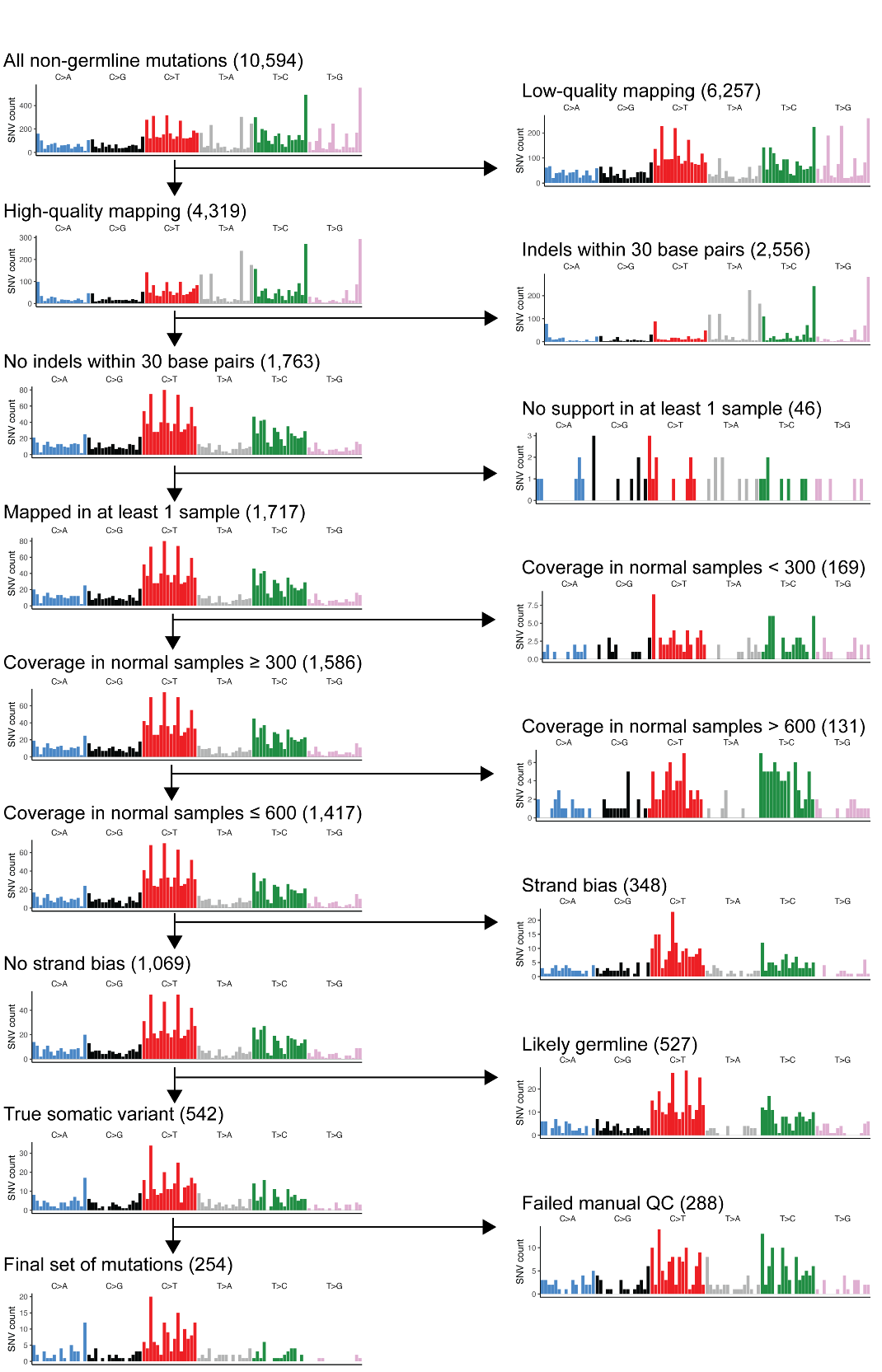

**Supplementary Figure 1. Filtering steps to obtain a set of high-quality somatic mutations.** The top plot shows all mutations (n = 10,954) retained after initial filtering or germline mutations. From top to bottom, we show trinucleotide contexts of mutations retained after each subsequent filtering step. To the right, we show trinucleotide context spectra of mutations removed after each filtering step. Numbers in brackets indicate the number of mutations in a given set.

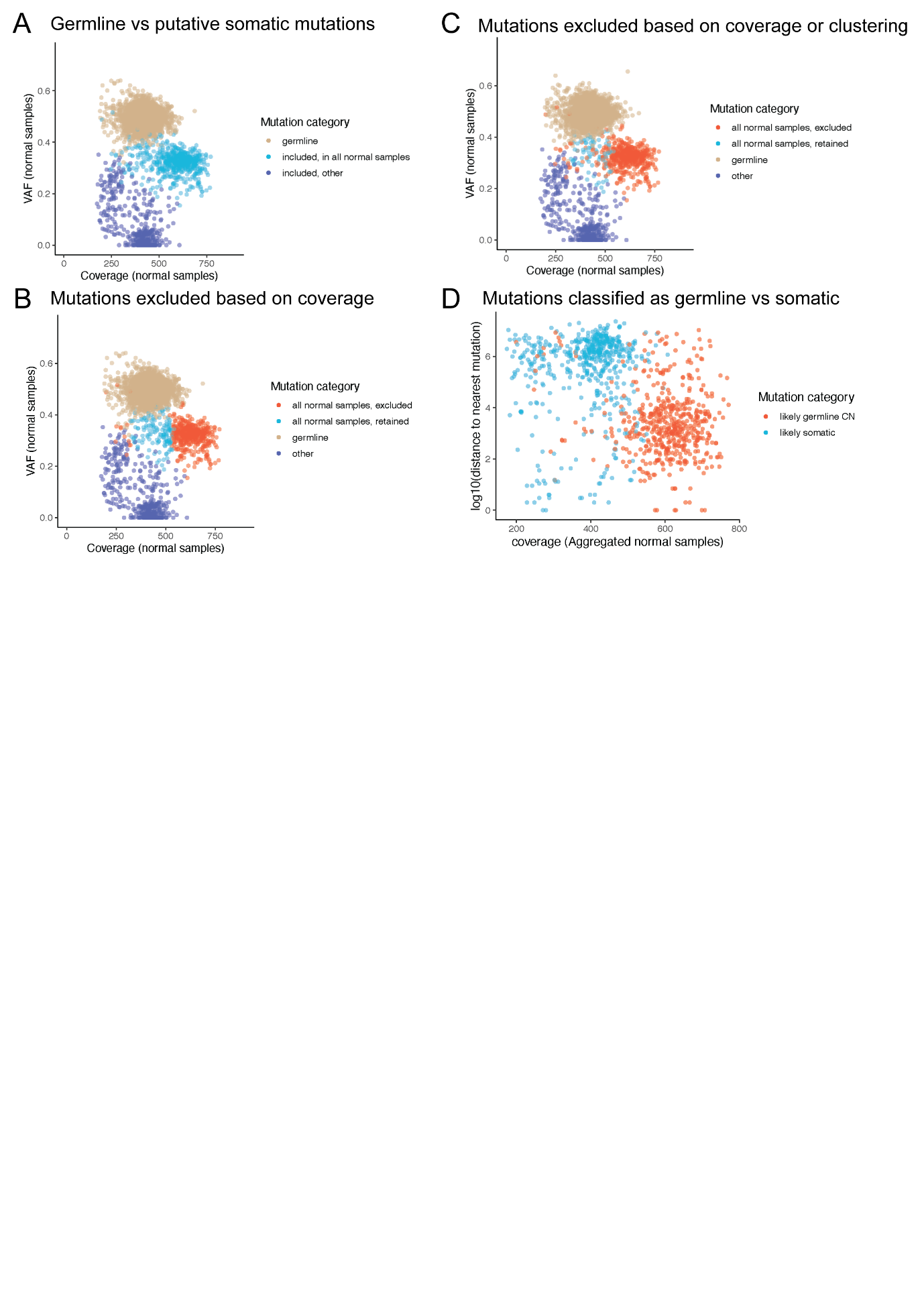

**Supplementary Figure 2. Filtering out germline mutations present on regions of altered copy number in the germline).** A. Clusters of mutations identified as germline from binomial test (‘germline’), retained mutations present in all normal samples and retained mutations not identified in all normal samples. Mutations present in all normal samples are present in a cluster that exhibits a higher coverage than other classes of mutations, suggesting they could be associated with copy number changes in the germline. B. Labelled mutations excluded following filtering out mutations with coverage in aggregated normal samples > 1.25x median or < 0.75x median of aggregated coverage in normal samples for all mutations identified on a given chromosome. C. Labelled mutations excluded following filtering out mutations with coverage in aggregated normal samples > 1.25x median or < 0.75x median of aggregated coverage in normal samples for all mutations identified on a given chromosome, or present in clusters of > 10 mutations in 50kb region. D. Validation of filtering. We plotted all 1,069 mutations retained in the analysis following application of the first filters and plotted their coverage in aggregated normal samples against the distance to the nearest mutation (log10). 542 mutations labelled as likely germline show lower distance to the nearest mutation and differences in coverage.

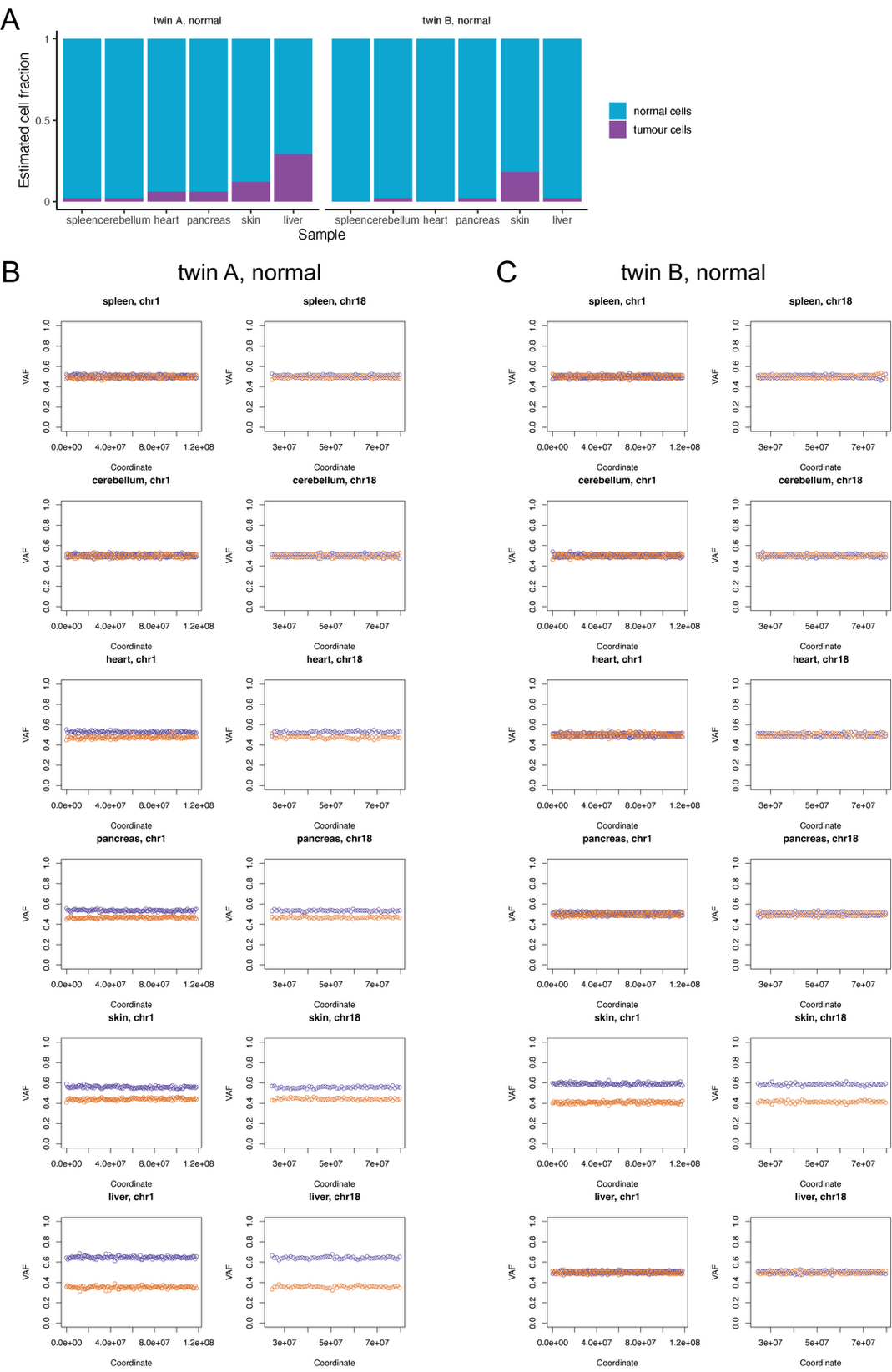

**Supplementary Figure 3. Estimates of tumour cell fraction in normal and tumour samples based on copy number alterations. Panel A** displays estimates of purity in each normal sample. **Panels B and C** present VAF of chromosome 1p and 18q in each normal sample from twin A (B) or twin B (C). Only the portion of the chromosome 1 or 18 which was lost in the posterior fossa tumour sample from twin A is shown, as we were only able to phase heterozygous SNPs located on this chromosome segment. Each point shows the median VAF of mutations present on either chromosome in a bin of 1 Mb.

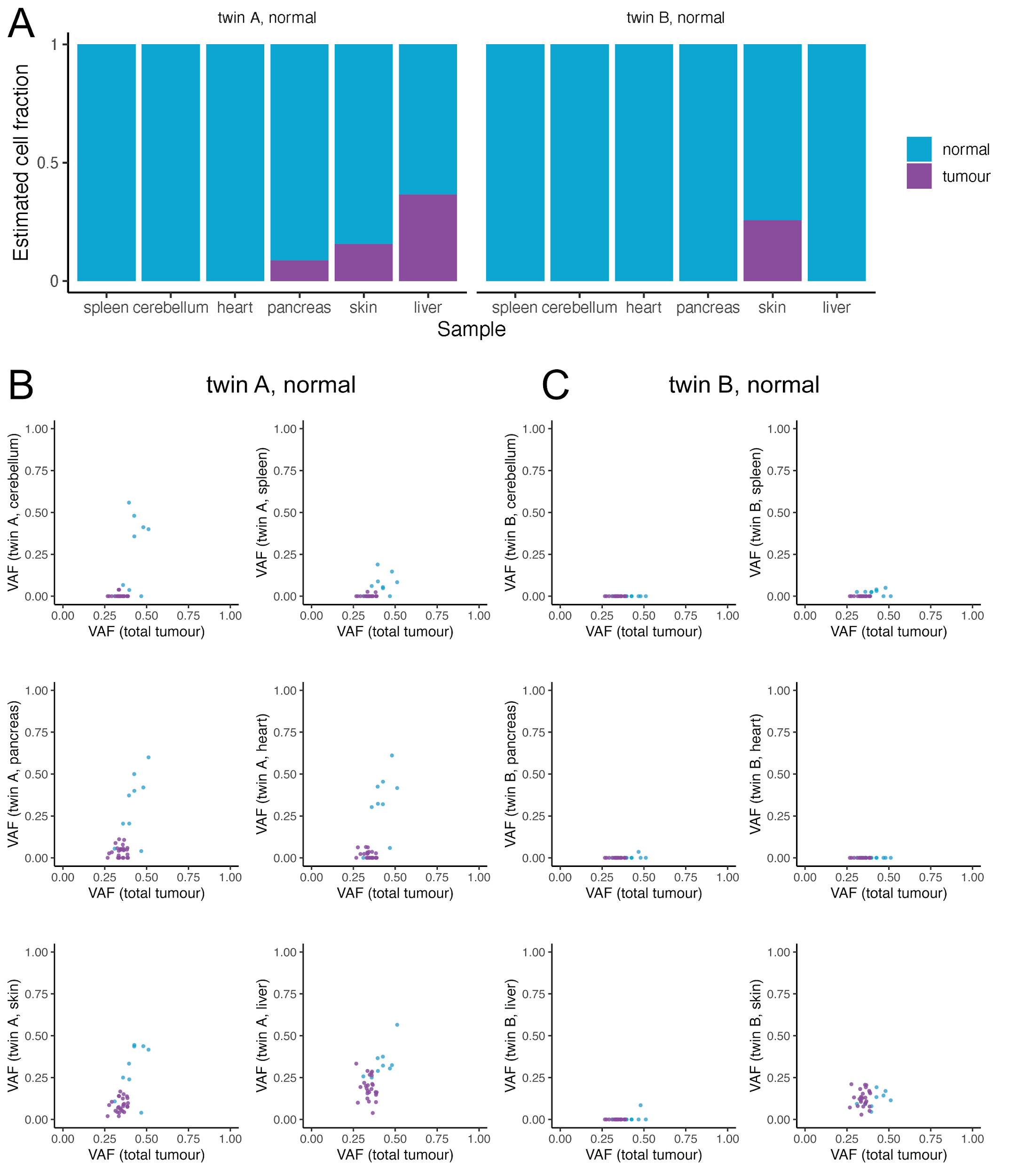

**Supplementary Figure 4. Estimates of tumour cell fraction in normal and tumour samples based on VAFs of early embryonic vs tumour-specific mutations. Panel A** Panel B shows estimates of the content of normal and tumour cells in each sample, based on the median VAF of 26 mutations classified as tumour-specific in this sample. C-D. VAF of 34 mutations in normal samples from twin A (C) and twin B (D). Mutations labelled as tumour specific or shared embryonic based on VAF in normal samples. In normal samples not infiltrated by tumour cells, VAF of tumour-specific mutations is 0.

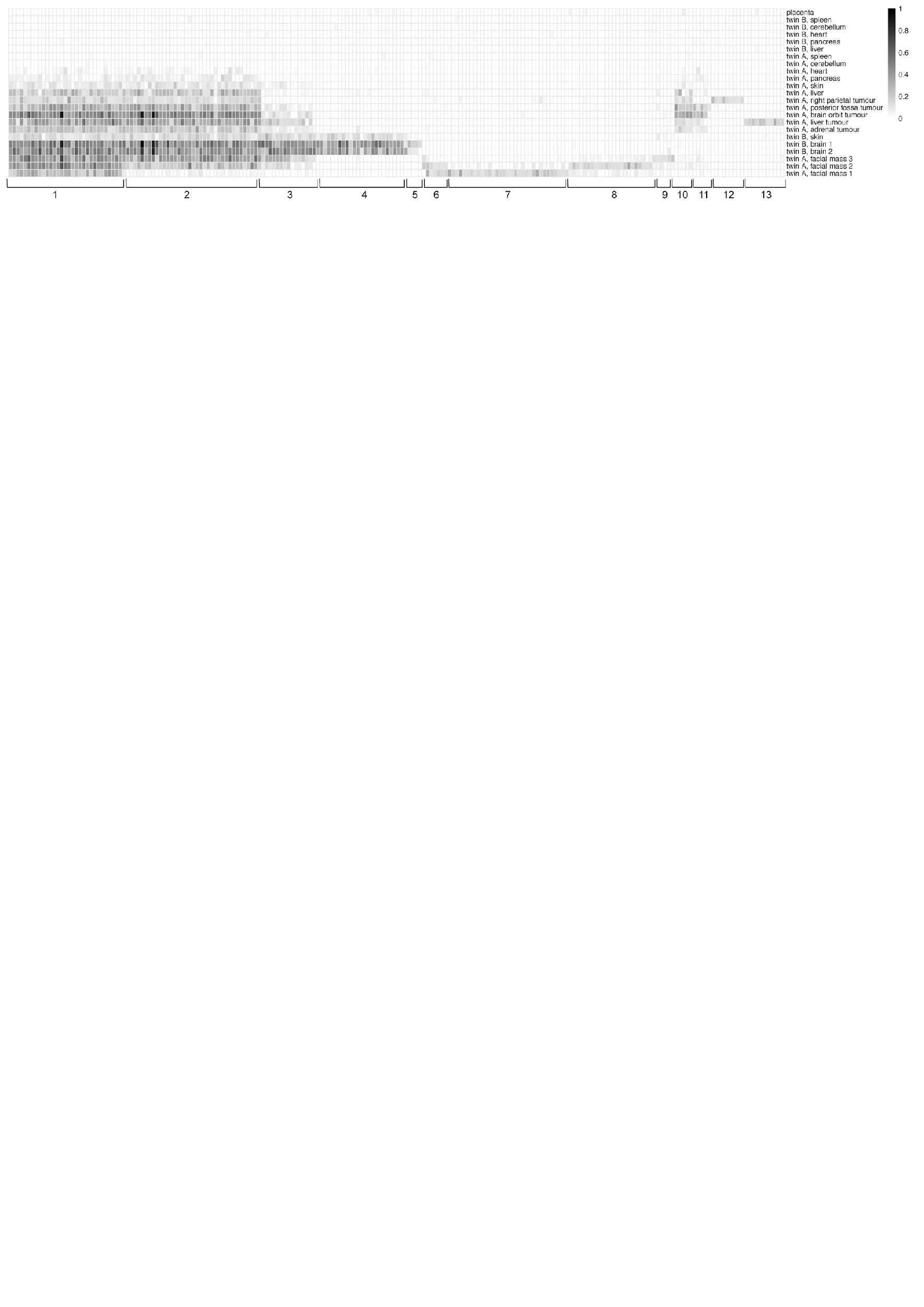

**Supplementary Figure 5.** Clustering of tumour specific mutations. Each vertical column represents one mutation. The colour scale represents VAF. Cluster labels 1-13 correspond to numbering on Figure 3C.

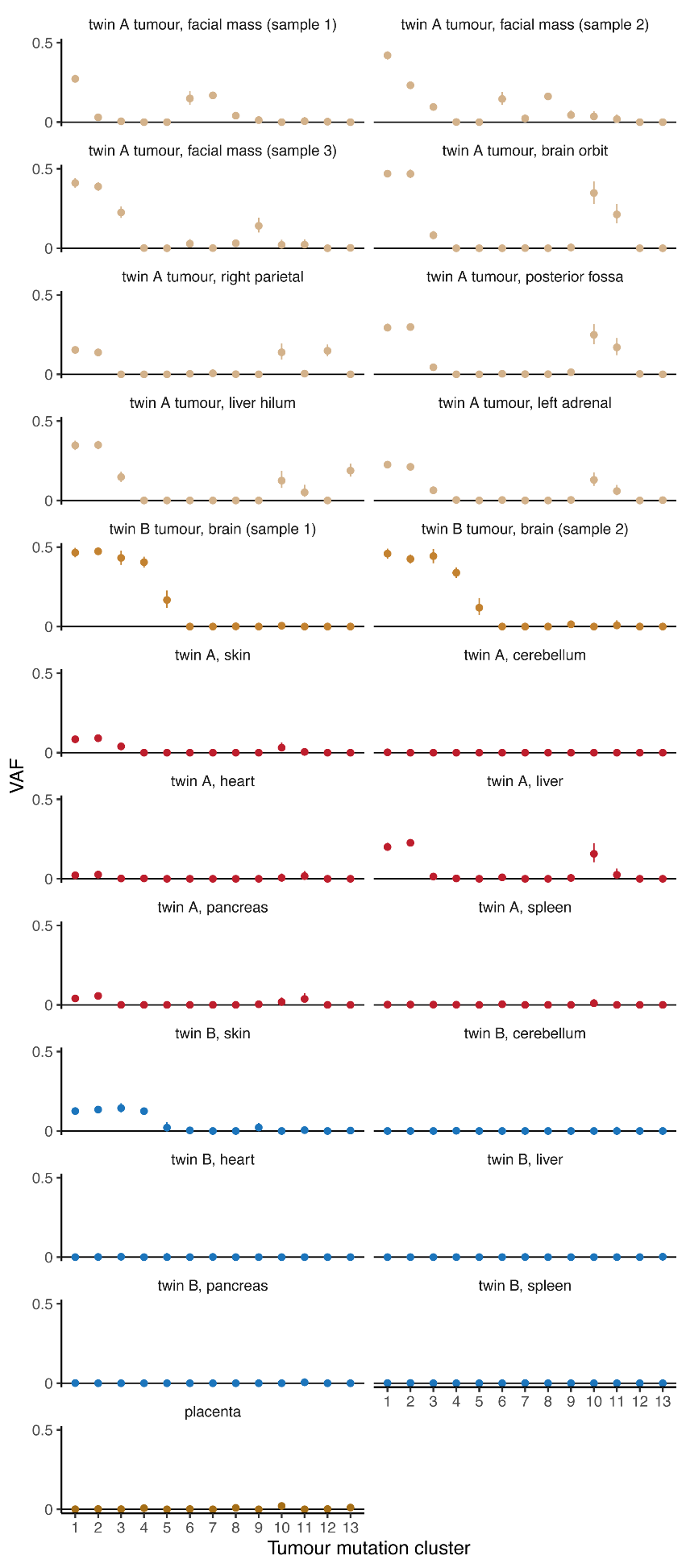

**Supplementary Figure 6**. Mean VAFs of each cluster of tumour specific mutation in each normal and tumour sample of both twins and the placenta (bulk)

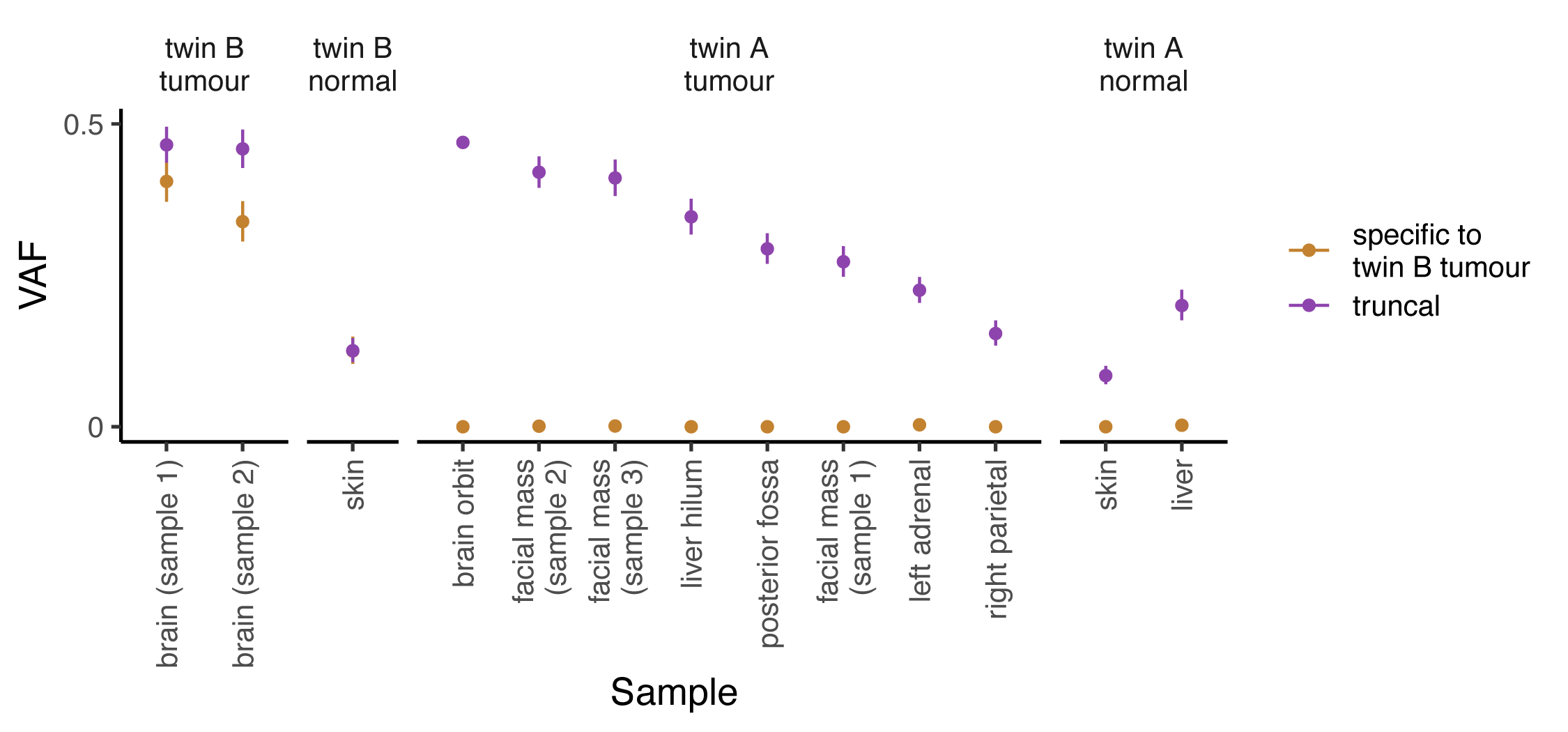

**Supplementary Figure 7.** Comparison of mutations in cluster 4 (specific to twin B tumour) and cluster 1 (truncal) in twin A and twin B samples.

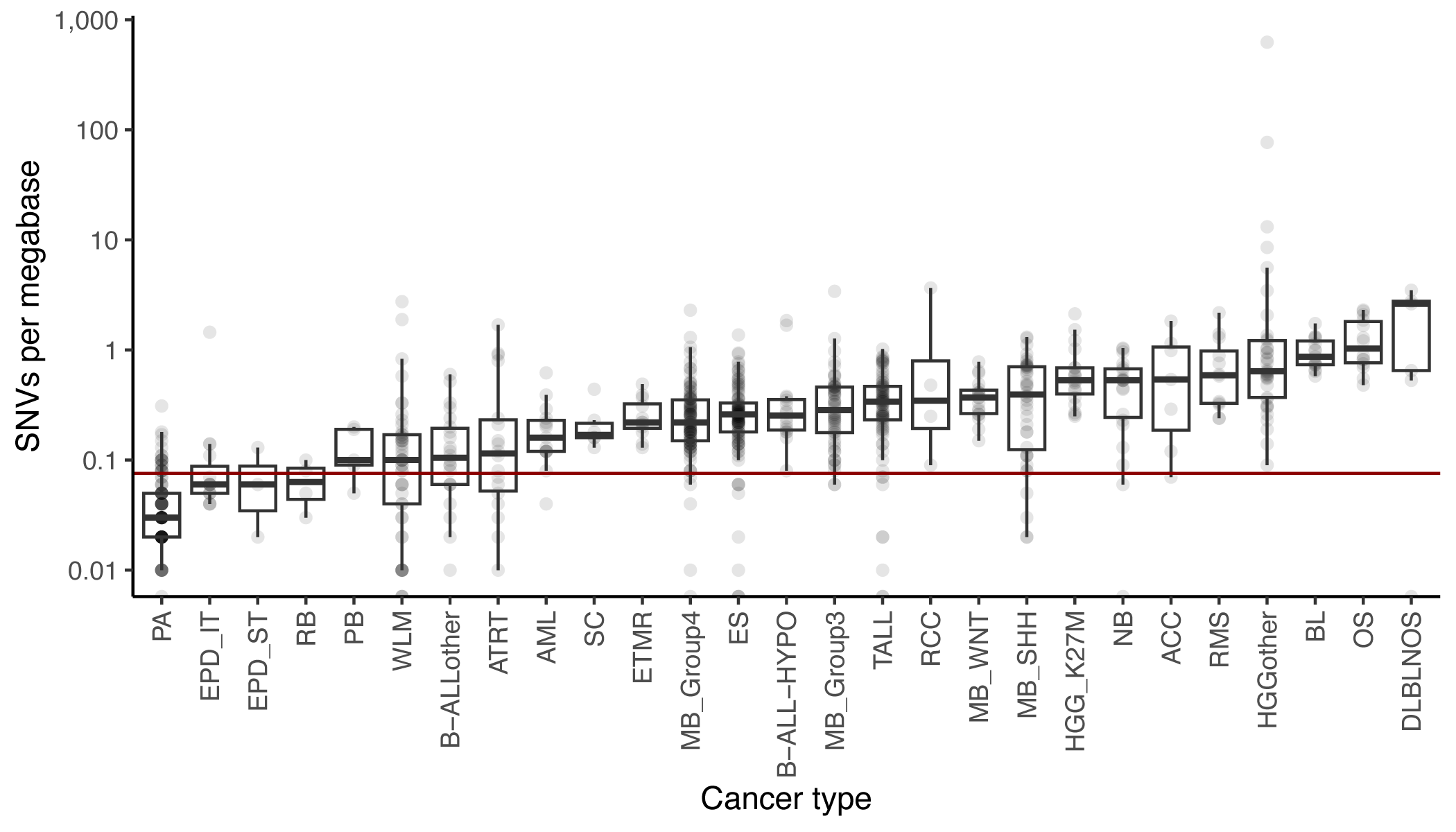

**Supplementary Figure 8. Comparison of tumour mutation burden.** The red line indicates the mutation burden observed in the tumour reported here. SNV – single nucleotide variant. Data for comparison are from Thatikonda et al.^13^

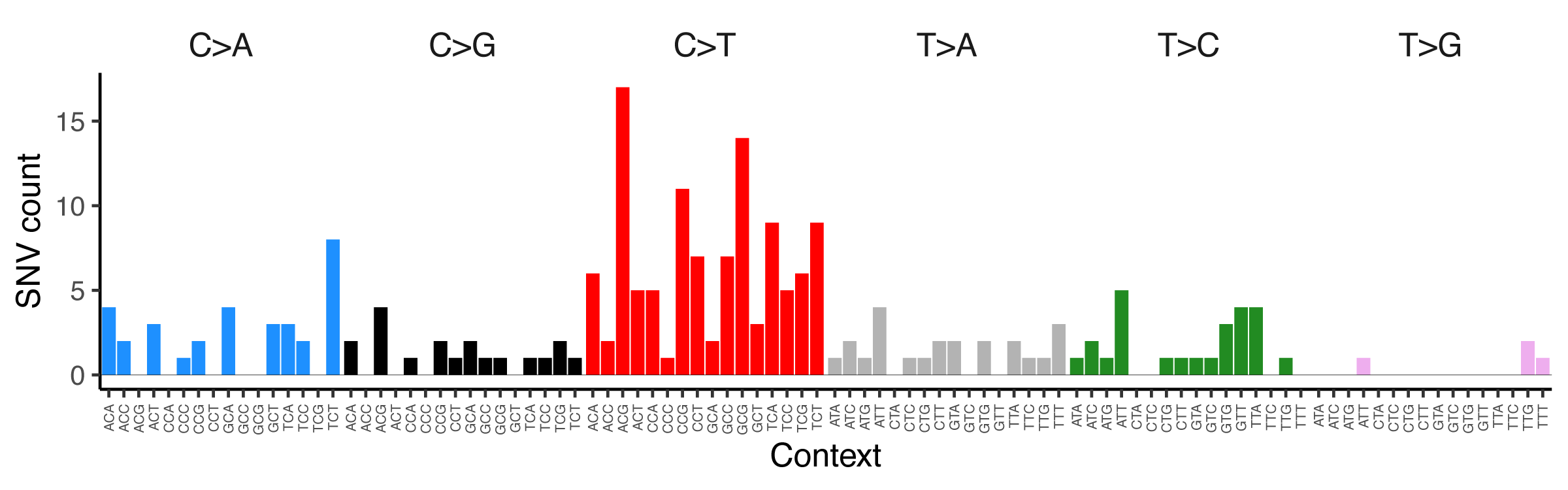

**Supplementary Figure 9**. Trinucleotide mutation spectrum of 212 tumour-specific mutations.

**Method and supplementary note references**
